# Tree diversity and density affect damage caused by the invasive pest *Cameraria ohridella* in urban areas

**DOI:** 10.1101/2022.04.30.490133

**Authors:** Alex Stemmelen, Hervé Jactel, Bastien Castagneyrol

## Abstract

Invasive, non-native invasive pests pose a growing threat to urban trees and the services they provide to urban residents. With the reluctance to use chemical insecticides in cities, environmentally friendly methods of pest management are needed. Tree diversity is known to affect insect herbivory in forest, with higher tree species diversity leading to lower level of damage. However, the validity of those findings for a non-native insect in an urban environment remains to be demonstrated.

We monitored 54 horse chestnut trees attacked by the invasive horse chestnut leafminer *Cameraria ohridella* in the city of Bordeaux, France. We analyzed the effects of neighboring tree diversity and density on the abundance, damage and parasitism rate of these leafminers.

We showed that the abundance and damage of *C. ohridella* significantly increased with higher local tree canopy cover. We found that the parasitism rate of *C. ohridella* increased with the species diversity of neighboring trees. However, this increase in parasitism rate was not associated with a decrease in leaf area damaged.

Our results pave the way for the management of exotic insect pests in cities based on the manipulation of spatial distribution and species diversity of urban trees.

## Introduction

Invasive insect pests represent a growing threat to forest health worldwide (Kenis et al., 2009; Venette & Hutchison, 2021), which has been exacerbated by increasing global trade and climate change (Roques, 2010). These biological invasions have a long lasting effect on the vitality and functioning of forest ecosystems and entail considerable economic costs (Zenni et al., 2021). Introduction of non-native forest insect pests is primarily driven by trading of live trees, imports of timber or use of wood packaging (Meurisse et al., 2019). As a result, nearly 90% of the first detections of non-native forest insects in Europe were made in urban forests, which are close to transportation hubs and concentrate high human populations (Branco et al., 2019). In cities, insect forest pests has been shown to compromise ecological and aesthetics values of trees and can pose a risk to the health of inhabitants, for example, through skin and respiratory irritation caused by the hairs of some herbivorous caterpillars (Tomlinson et al., 2015; Tooker & Hanks, 2000). Once established in an urban environment, these insects also benefit from favorable development conditions with higher temperatures that accelerate their development and water stresses that make the trees more vulnerable (Dale & Frank, 2017, Dale & Frank, 2018). Therefore, measures need to be taken to mitigate the damage caused by non-native forest pests in urban area and to better understand the ecological drivers likely to slow their spread before they escape out of cities and colonize nearby rural forests.

Due to their adverse impact on the environment (Mahmood et al., 2016) and human health (Blair et al., 2015; Margni et al., 2002), chemical insecticides are increasingly banned from urban areas worldwide (Kristoffersen et al., 2008) and priority is now given to preventive methods of pest management. In particular, many studies have shown that herbivorous insect are less abundant and inflict less damage on trees surrounded by heterospecific neighbors (Guyot et al., 2019; Jactel et al., 2021). This phenomenon, called associational resistance, can be explained by two main non-exclusive mechanisms. First, the presence of heterospecific neighboring trees may limit the ability of an herbivorous insect to locate its host tree (the resource concentration hypothesis, Hambäck & Englund, 2005; Root, 1973). Second, higher tree diversity can favor herbivore’s natural enemies (Stemmelen et al., 2022), by increasing the availability of suitable habitats or alternative resources such as prey for predators, pollen or nectar for parasitoids, leading to a better control of insect pest population (the natural enemies hypothesis, Risch et al., 1983; Root, 1973). However, while associational resistance and the hypothesis mentioned above have received large support in forests ecosystems (Field et al., 2020; Poeydebat et al., 2021; Tudoran et al., 2021; Jactel et al., 2021), evidence of its validity in urban environments remain scarce. Urban environments greatly differ from natural forest environments. Urban trees are often more patchily distributed and urban forest patches are often separated by large hostile non-forest habitats. Thus, the high fragmentation of urban forests could limit colonization capacities of herbivorous insect or result in increased insect mortality during the prolonged dispersal phase required to locate isolated hosts trees, which could eventually reduce insect herbivory. An alternate, although non-exclusive prediction, is that insect herbivores, once established on the fewer available host trees, would limit their dispersal and keep damaging the same isolated host trees, thus increasing local damage.

*Cameraria ohridella* Deschka & Dimić is an invasive forest pest from the Gracillariidae family firstly discovered in Macedonia in 1984 (Deschka & Dimić, 1986). It rapidly invaded all Europe and mainly feeding on horse-chestnut *Aesculus hippocastaneum* L., an important ornamental tree in many European cities. The larvae mine the leaves, creating brown lesions that can eventually cover the entire leaf area and cause early defoliation. Although mature trees are rarely endangered by such defoliations, they can result in serious loss in radial growth and aesthetic values (Jagie∤∤o et al., 2019; Myśkow et al., 2021), while mortality may occur in highly infested young trees (Salleo et al., 2003). Past studies in urban areas have shown that *C. ohridella* can be parasitized by insects (Girardoz, Quicke, et al., 2007a; Grabenweger et al., 2010; Volter & Kenis, 2013) and preyed by birds (Grabenweger, Kehrli, et al., 2005b). Parasitoids are the most studied natural enemies of *C. ohridella*, with more than 30 indigenous species found in Europe (see https://www.cabi.org/isc/datasheet/40598). Parasitism rate usually varies between 1% and 20% (Freise et al., 2002; Girardoz, Quicke, et al., 2007a; Grabenweger, 2003; Grabenweger et al., 2010) and occurs mainly on the nymph stage. It has been also reported that some parasitoids (e.g. *Baryscapus nigroviolaceus, Cirrospilus elegantissims)* parasitizing leaf miners of the sycamore maple *Acer pseudoplatanus* can shift to *C. ohridella* (Girardoz et al., 2007b). *C. ohridella* is now widely distributed in Europe and has been present for many years, which may have increased the likelihood to recruit local enemies that were generalist enough to switch on this new host.

In this study, our goal was to assess the role of neighbouring trees on the abundance of the non-native insect pest *C. ohridella*, its damage to the host *A. hippocastaneum*, and the top-down control by natural enemies in an urban environment. To address this, we measured leaf area damaged by C. orhidella larvae and abundance of *C. orhidella* flying adults on 54 horse chestnuts of the city of Bordeaux, France. We concomitantly assessed the parasitism of *C. orhidella* pupae and linked levels of herbivory and parasitism to tree neighborhood variables, namely density and diversity. In particular, we predicted that leaf damage would be lower in more diverse tree neighborhood, in accordance to the resource concentration and natural enemies hypotheses. Additionally, we predicted that herbivory would decrease with increasing tree density and number of conspecific horse chestnut in the neighborhood, following the dilution of damage when host density is higher. Finally, we expected that both tree diversity and quantity of maple trees in the neighborhood of focal chestnuts would increase the parasitism rate by parasitoids, resulting in lower leaf damage by *C. ohridella*. In doing so, our study seeks to advance knowledge of the ecological drivers of urban tree resistance to invasive pests and provides insights into the management of their populations in urban forests.

## Material & Methods

### Study system

The study was conducted in the city of Bordeaux (France, 44°50’N, 0°34’W), with an average temperature of 13.8°C and average annual rainfall of 803 mm. Horse-chestnut is one of the most common tree species in the city, being the 10^th^ most planted in the public domain. *Cameraria orhridella* has three generations per year in the study area. Adults emerge in May from nymphs overwintering in dead leaves. Larval development lasts for 25-35 days and passes through four (occasionally five) feeding instars and two spinning instars (Skuhravý, 1998). Specialized mouthparts allow the larvae to mine the leaf parenchyma. Then larvae go through two spinning instars. Adults of the first generation emerge in June, while adults of the second and third generations emerge in early August and in September. Depending on the weather conditions, a fourth generation can sometimes occur. A fraction of insects of each generation overwinters as nymphs on dead leaves and emerge the next year to (re)colonize horse chestnut trees (Fig. 1).

**Figure 1.**
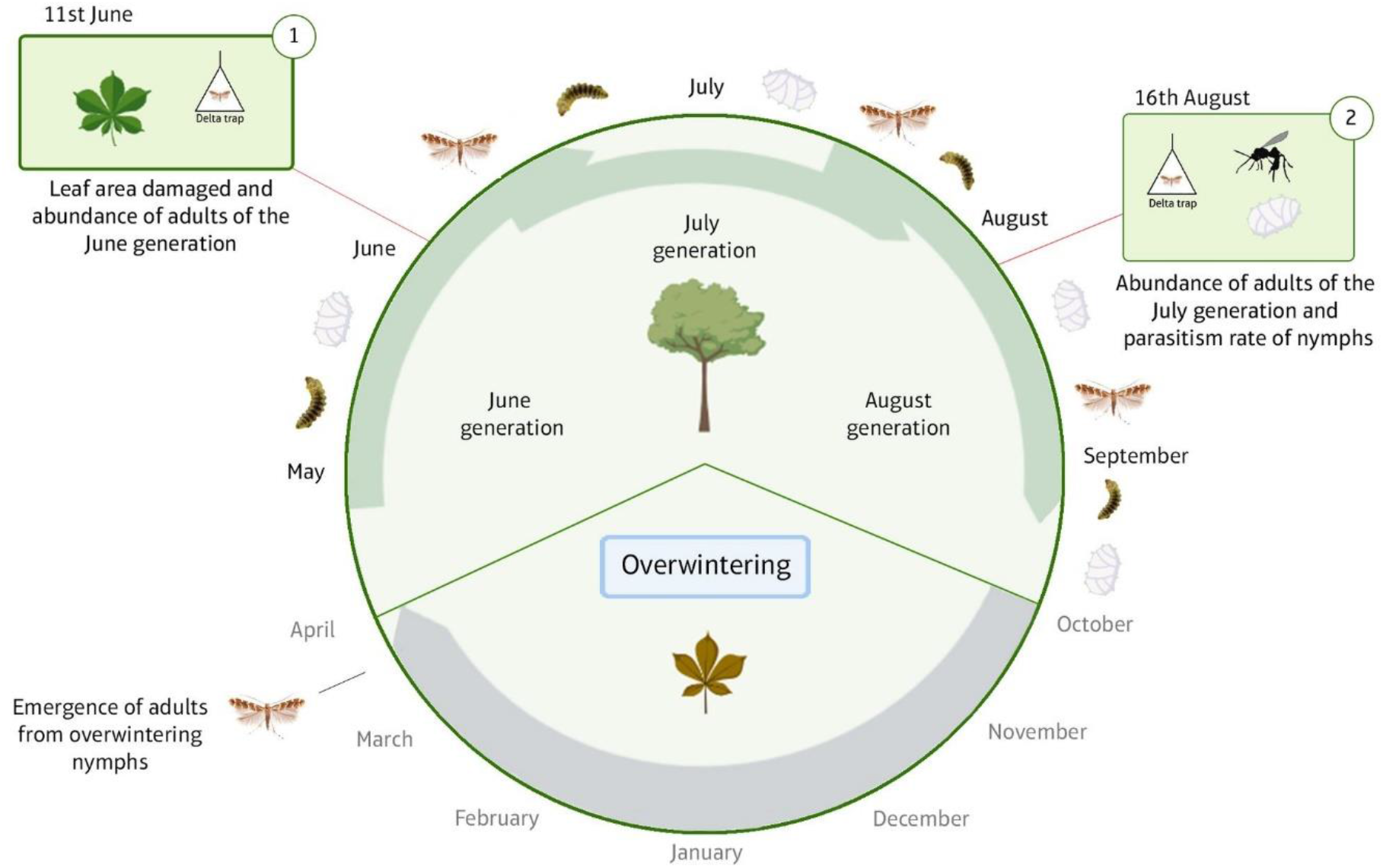
Simplified life cycle of *C. ohridella* in Bordeaux, from the emergence of adults and the development of each generation in the canopy of horse chestnut in late March to the overwintering in leaf litter in October. The two green frames indicate when (1, June) leaves were sampled for leaf damage assessment and adults of the 1st generation were trapped and (2, August) adults of the 2^nd^ generation were trapped and chrysalis collected for parasitism rate assessment.

### Study site and tree selection

In 2020, we estimated canopy cover in a 100m buffer, and tree diversity using tree species richness and Shannon diversity index in a 20m buffer around the 772 horse chestnuts of the public domain of Bordeaux. We then made a selection of horse chestnuts based on the following three criteria: (1) lack of significant correlation between tree diversity and tree density in the buffers, (2) lack of spatial autocorrelation and (3) largest distance between selected trees *(minimum pairwise distance: 2m; max*.: *7240m; mean: 1903m*). The final tree selection consisted in 54 horse chestnuts distributed along two orthogonal gradients of tree density and diversity (Supplementary material).

### Tree neighborhood

Every tree of the public domain of Bordeaux is geolocalized, identified to the species level and has its age and height referenced. This information is available through the city database for urban trees (https://opendata.bordeaux-metropole.fr/explore/dataset/bor_arbres/information/). We described tree neighborhood around each focal horse chestnut by assessing tree density, tree diversity, number of conspecific horse-chestnuts and number of maple trees. Canopy cover was assessed in buffer of 100m. Tree diversity was assessed in buffer of 20m around each focal tree, both using tree species richness and Shannon diversity index. Number of conspecific horse chestnut and number of maple were assessed in buffer of 100m. Buffer sizes were chosen in regard to *C. ohridella* poor flight dispersal capacity (Augustin et al., 2009) and in order to get a precise estimation of tree diversity in the neighborhood. We verified *a posteriori* that no trees in private properties – which were not included in the data base – were included in the 20m buffers. These variables were assessed using Geographical Information System (QGIS Development Team, 2018) and R software version 3.4.4 (R Core Team 2019).

### Abundance of *C. ohridella* and associated damage

In June 2020, we haphazardly collected 30 leaves on each sample trees. The leaves were collected with a pole pruner by rotating around the tree so as not to favor one aspect over another. We preserved the leaves in paper bags and brought them back to the lab for herbivory estimation. We estimated the damage caused by the horse-chestnut leafminer by evaluating the leaf area covered by mines. We took pictures of each leaves and processed them using the Inkscape software. Mines were manually cropped on the images and the percentage of leaf area damaged was calculated as the ratio between the area covered by mines and the total leaf area. Mean leaf area damaged was calculated across the 30 leaves collected on each tree.

Additionally, we conducted two surveys to estimate the number of flying adults of the first and second generation around each sample horse chestnut. We used delta sticky traps combined with the sex pheromone of *C. ohridella* (Pherobank B.V.) to capture and count flying adults. Traps were hung to low branches of chestnut trees and left in place for 6 days in late spring (First survey, from June 11^st^ to June 16^th^, 2020) and for 8 days in summer (Second survey, from August 18^th^ to August 25^th^, 2020, Fig. 1). Traps were assessed once at the end of each survey and adults were counted by a single observer (AS).

### Parasitism by natural enemies

In August 2020, we collected 30 additional leaves with mines visible from the ground on each sample tree. Attacked leaves were sealed in plastic bags and brought back to the lab. We dissected the mines under a binocular magnifier, extracted the nymphs and placed them in Petri dishes. We then waited a full year for the parasitoids to hatch from the nymphs and counted the rate of parasitism across all leaves (Fig. 1). A single observer (AS) conducted nymphs extraction and parasitoid sighting.

### Statistical analyses

We used model selection coupled with model averaging to identify the best model fitting our data and used it to estimate model coefficient parameters (Grueber, Nakagawa, Laws & Jamieson, 2011). We first built a full model using all explanatory variables of interest (predictors). We excluded all sub-models that included correlated predictors and we standardized explanatory variables using Gelman’s approach (Gelman, 2008) to ease interpretation of parameter estimates after modal averaging. We then applied a procedure of model selection by running every model nested within the full model. We used Akaike’s criterion corrected for small sample size (AICc) as information criterion. We ranked all models based on difference in AICc between each model and the top ranked model with the lowest AICc (ΔAICc). When multiple models had a ΔAICc < 2, we used a model averaging approach to build an average model including all variables found in the set of best models. A given predictor was considered significant if the 95% confidence interval of its parameter estimate did not overlap zero. Finally, we assessed the relative importance of each parameter retained in models. The relative importance is a measure of the prevalence of each parameter in each model used during the model-averaging process. We used this approach three times, for the following three models:

#### Model 1: Mean leaf area damaged

We used a linear model to explain the variability in mean leaf area covered by mines using tree neighborhood variables and parasitism rate. The full model included the following explanatory variables: Tree canopy cover in 100m buffer *(Canopy)*, tree diversity *(Shannon* and *Richness)*, number of maples *(Acer density)* and horse chestnuts *(Aesculus density)* in a 100m buffer and parasitism rate *(Parasitism)*. It is important to note that parasitism rate has been assessed in August while mean leaf area damaged has been assessed in June. However, it is likely that tree with a high level of parasitism in August had similar level of parasitism in June, making the inclusion of parasitism rate in this model relevant.

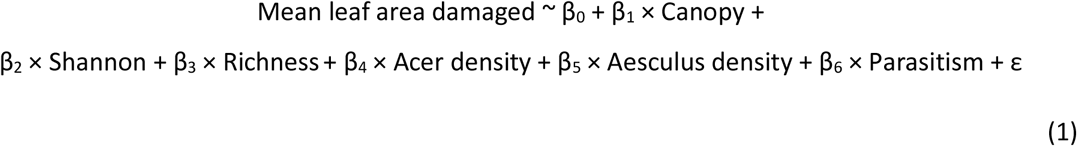

Where *β* are model coefficient parameters and *ε* the residuals. Richness and Shannon were auto-correlated and as such, did not appear together in sub-models.

### Model 2: Abundance of *C*. *orhridella* adults of the first generation

We modeled the abundance of adults as a function of tree neighborhood density and diversity, while using the damage made by the larval stage as a covariate. By doing this, we can control that the adults captured come mainly from the same tree on which damage has been caused. We used a generalized linear model with a Gamma distribution family. The full model included the following explanatory variables: Tree canopy cover in 100m buffer *(Canopy)*, tree diversity *(Shannon* and *Richness)*, number of maples *(Acer density)* and horse chestnuts *(Aesculus density)* in a 100m buffer and mean leaf area damaged *(Leaf area damaged)*.

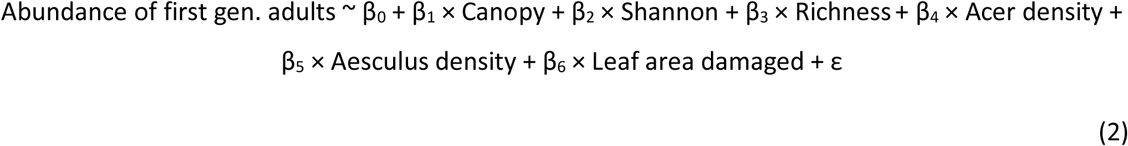

#### Model 3: Parasitism rate

We tested the effect of tree neighborhood density and diversity on parasitism rate while controlling for potential density-dependent effects by accounting for the abundance of adults of the previous generation. In 12 trees, leaf-miner infestation rate was so low that we could not sample 30 leaves with visible damage by mines. We only included in this analysis trees from which we could extract 5 nymphs or more (n = 42). As the number of nymphs parasitized did not follow a Normal or Poisson distribution, we chose to treat it as binomial variable (0 = no pupae parasitized; 1 = at least one pupae parasitized). We used a generalized linear model with a binomial distribution family. The full model included the following explanatory variables: Tree canopy cover in 100m buffer *(Canopy)*, tree diversity *(Shannon* and *Richness)*, number of maples *(Acer density)* and horse chestnuts *(Aesculus density)* in a 100m buffer and abundance of flying adults of the second generation *(Second gen. adults)*.

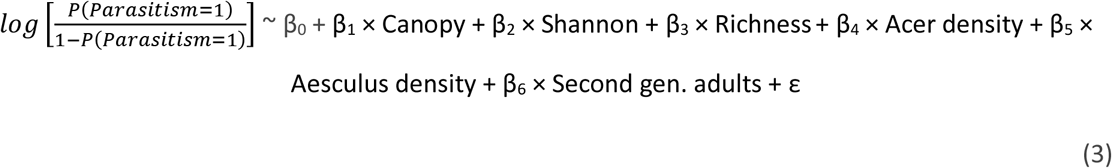

Statistical analyses were performed using the R software version 3.4.4 with packages lme4 (Bates, Mächler, Bolker, & Walker, 2015) and MuMIn (Barton 2019).

## Results

Canopy cover ranged from 7.20% to 68.44% of the total area in 100m buffers around focal chestnut trees, with a mean ± SE of 37.38 ± 2.52%. Tree species richness in a 20m buffer was on average 4.53 ± 0.41 and ranged from 1 to 13 tree species while Shannon diversity of trees ranged from 0 to 2.44, with a mean of 1.13 ± 0.098. Number of neighboring horse chestnut in a 100m ranged from 1 to 43 with a mean of 12.33 ± 1.40 and number of neighboring maple in a 100m ranged from 0 to 17 with a mean of 2.29 ± 0.56.

### Abundance of *C. ohridella* and associated damage

On average, pheromones traps caught 572.6 ± 89.8 (median: 228) flying adults of the first generation and ranged from 23 to 2838 adults caught. They caught 87.5 ± 14.2 (median: 46) adults of the second generation. We observed the observed the presence of *C. ohridella* in 48 (89%) of the the 54 trees selected. Mines of horse-chestnut leaf-miner covered on average 5.3 ± 1.0 % of the leaf area. Among trees with mines, the percentage of leaf area covered by mines varied from 0.014% to 34.6%.

Mean leaf area damaged by mines significantly increased with the canopy cover in a 200m buffer around focal trees (average model coefficient parameter estimate ± CI : 8.9 ± [4.8; 12.9], n = 54, Fig. 2, Table 1, model 1).

**Table 1.** Summary of model coefficients estimated through model averaging of the models with ΔAICc< 2. Bold characters indicate that parameters are significant. Estimate and SE of the model 2 are based on a generalized linear model with a Gamma distribution family.

| Models |  |  |  |  |  |
| --- | --- | --- | --- | --- | --- |
| Parameter | Estimate | Adjusted SE | 95% CI | Relative importance | P value |
| **Model 1 - Leaf area damage** |  |  |  |  |  |
| **(Intercept)** | **5.560** | **1.00** | **(3.58; 7.54)** |  | **< 0.001** |
| **Canopy cover** | **6.950** | **2.15** | **(2.74; 11.17)** | **1** | **< 0.001** |
| Acer density | 0.360 | 1.20 | (-2.69; 6.30) | 0.3 | 0.716 |
| **Model 2 - Abundance 1st generation adults <sup>1</sup>** |  |  |  |  |  |
| **(Intercept)** | **2.910** | **0.49** | **(1.95; 3.88)** |  | **< 0.001** |
| **Canopy cover** | **-2.770** | **0.78** | **(-4.3; -1.24)** | **1** | **< 0.001** |
| **Leaf area damage** | **-0.570** | **0.24** | **(-1.05; -0.097)** | **1** | **0.018** |
| Aesculus density | -0.140 | 0.54 | (-2.57; 1.25) | 0.23 | 0.78 |
| Richness | -0.069 | 0.29 | (-1.42; 0.77) | 0.21 | 0.81 |
| **Model 3 - Parasitism rate** |  |  |  |  |  |
| **(Intercept)** | **-0.440** | **0.31** | **(-1.04; 0.16)** |  | **0.154** |
| **Shannon** | **1.610** | **0.67** | **(0.27; 2.94)** | **1** | **0.017** |
| Canopy cover | 0.150 | 0.41 | (-0.69; 1.85) | 0.26 | 0.716 |
| Acer density | 0.080 | 0.32 | (-0.79; 1.58) | 0.21 | 0.794 |

<sup>1</sup> Estimate, SE and 95% CI value have been multiplied by 1000.

**Figure 2.**
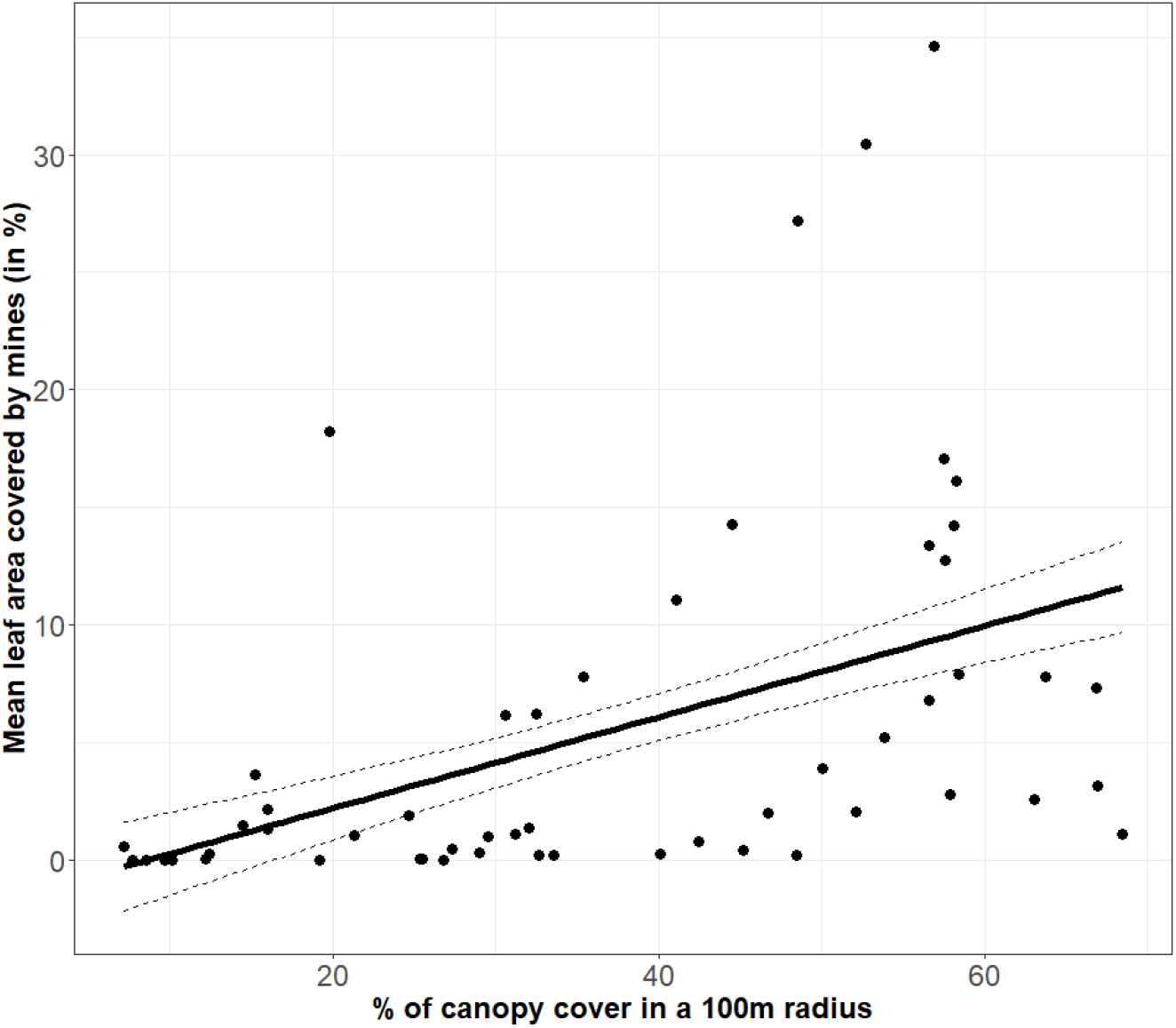
Effects of canopy cover in a 100m radius on mean leaf area covered by mines. Dots represent individual trees. Solid and dashed lines represent prediction and adjusted standard error of the average model (Table 1).

Once leaf area damaged was accounted for in the model, the abundance of first generation flying adults was correlated with tree cover in a 100 m buffer zone around each focal tree (−2.79 ± [-4.3; - 1.24], n = 54, Table 1, model 2). Leaf area damaged was correlated with abundance of adults of the second generation (−0.57 ± [-1.05; -0.09], n = 54, Table 1, model 2).

### Parasitism rate

Of the 909 pupae extracted from the mines of *C. ohridella*, 49 were parasitized (5.4%). Parasitized chrysalises were found in 22 horse chestnut trees (out of 42, *i*.*e*., 40.7%). Five models had a ΔAICc < 2 in comparison to the top ranked model (Supplementary material 2); they included Shannon diversity, canopy cover, tree height and number of neighboring maple as predictors. The probability of a pupae being parasitized significantly increased with increasing tree diversity in a 20m buffer (1.6 ± [0.2; 3.0] n = 42, Fig. 3, Table 1, model 3). The coefficients of other predictors retained in the range of models with ΔAICc < 2 were not statistically different from zero.

**Figure 3.**
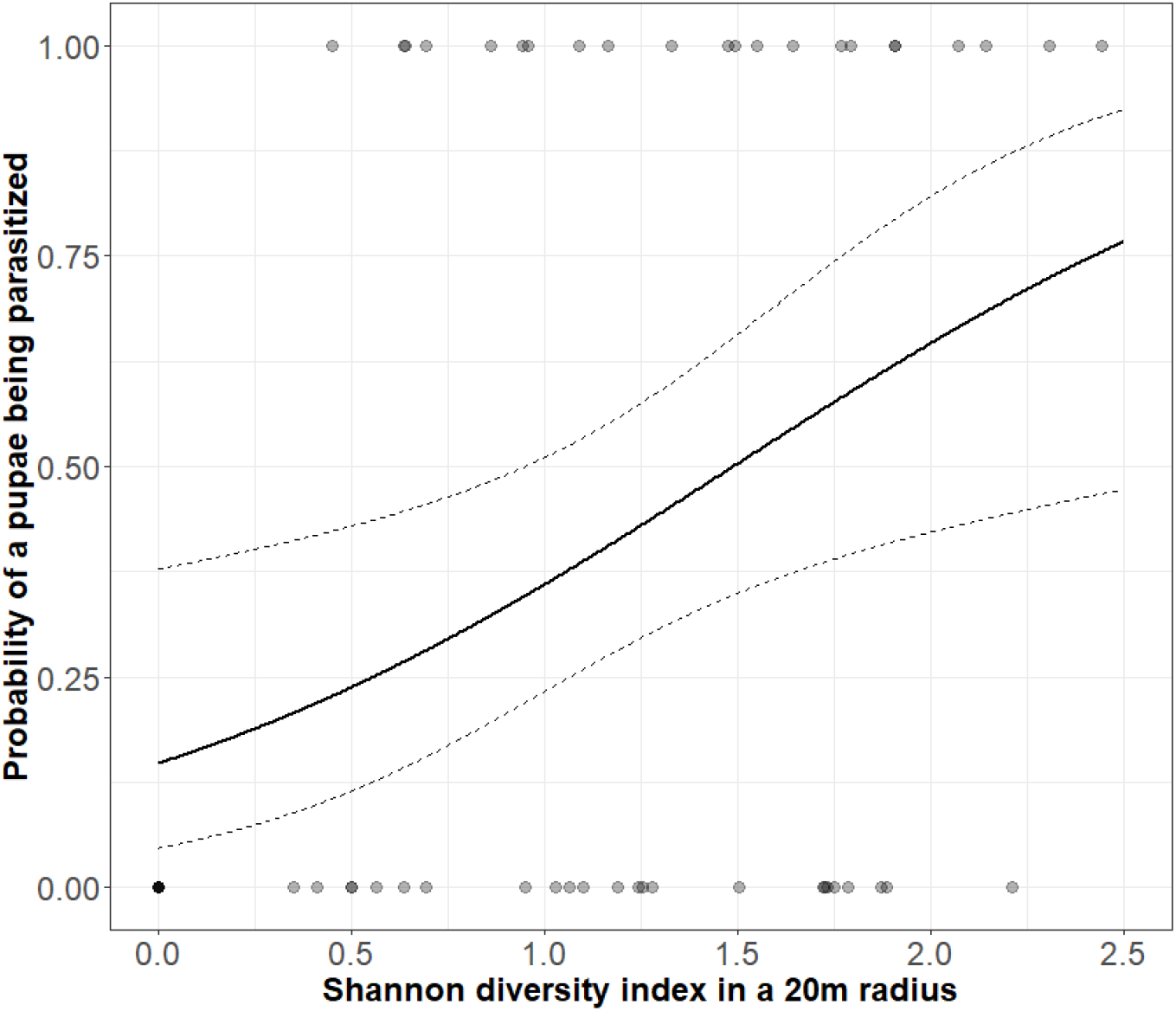
Effects of tree diversity on the probability of a pupa to be parasitized. Solid and dashed lines represent prediction and adjusted standard error of the average model (Table 1).

## Discussion

We showed that tree neighborhood can influence insect herbivory by an invasive tree pest in urban environments. Specifically, tree canopy cover, a proxy of tree density, had a significant and positive effect on the abundance of *C. ohridella* and associated damage. Additionally, we confirmed that tree diversity had a significant and positive effect on the parasitism rate by natural enemies of *C. ohridella*. Although this relationship was not associated with a decrease in leaf area damaged when tree diversity was high, our study demonstrates that some of the ecological mechanisms controlling insect damage in forest ecosystems can be transposed to urban areas where they can serve as basis for the pest management in urban forests.

Contrary to our predictions, *C. ohridella* damage did not increase with increasing number of conspecific horse chestnuts or decreasing tree diversity around the focal tree. This contradicts the resource concentration hypothesis that denser and monospecific stands of a host plant should suffer more herbivory damage (Grossman et al., 2018; Hambäck & Englund, 2005; Root, 1973). Even in the case of a specialist herbivore such as *C. ohridella*, increasing the resource concentration did not increase the amount of damage caused by the insect. The underlying processes of this theory is that herbivores would have a higher probability of locating and entering these patches with a high concentration of host plants but also a low propensity to leave these patches notably because of an abundant resource. Here we found that damage by *C. ohridella* increased with increasing canopy cover around a focal horse chestnut and this variable may reflect the primary mechanism conditioning herbivore fluxes in host tree patches. Horse chestnuts surrounded by high amount of canopy cover are often located in parks or urban green spaces, where tree density is important, while isolated trees are often found in highly urbanized space, such as parking lots or along main roads. It has been previously reported that one of the most efficient way of controlling the spread and damage of *C. ohridella* is the removing of dead leaves, in which nymphs overwintered until the following year (Kehrli & Bacher, 2003, 2004). In parking or alongside roads, dead leaves are often collected by public services or carried away by the passing of cars, which might impaired the ability of *C. ohridella* to recolonize each year and thus further prevent the damage caused to isolated horse chestnut. Conversely, public policies often go against leaf removal in parks and urban greens spaces, and the presence of understory vegetation can limit leaf dispersal by the wind. As a result, the infestation can reoccurs more easily each year, resulting in a higher level of damage in urban areas where trees are more abundant, irrespective of their tree species diversity. It is then likely that the specificity of the urban environment disrupted the effect of resource concentration on the recruitment of *C. ohridella*, a specialist pest, and associated damage.

Management practice may not be the only explanation behind the pattern we observed. Invasion process by non-native pest goes through three phases: arrival, establishment and spread (Liebhold & Bascompte, 2003; Williamson, 1996). During the establishment phase, isolated trees are less likely to host a great number of pests, as they are more difficult to locate, often leading to smaller population of insect compared to area with denser vegetation, especially in the case of a pest with poor dispersal capacity by flight (Gilbert et al., 2003; Valade et al., 2009). It has been previously theorized that smaller population are more vulnerable to demographic and environmental stochastic effects (Lande, 1993) which could lead to population extinction or loss of population’s fitness, namely the Allee effects (Fieberg & Ellner, 2001; Liebhold & Bascompte, 2003; Stephan & Wissel, 1994). Those density-dependent mechanisms such as failure to locate mates or inbreeding depression could help explain why abundance and damage of *C. ohridella* remained low in isolated trees where initial establishment phase led to small populations. Although *C. ohridella* is now well established in this area and the probability of this stochastic events occurring is reduced, taking better account of urban tree management practices as well as the demographic and dispersal processes of insect populations appears crucial to better understand the mechanisms conditioning the levels of damage caused by non-native pests in cities.

We found a positive effect of tree species diversity on the parasitism rate by natural enemies. Nymphs collected on horse chestnuts in more diverse environments were more often parasitized than in less diverse environments. This result provides support to the natural enemies’ hypothesis, which states that natural enemies are more abundant or active in more diverse plant communities and in particular in mixed forests (Stemmelen et al., 2022). Although urban studies often found little to no effect of vegetation complexity and diversity on natural enemies (Rocha et al., 2018; Sattler et al., 2010), some detected positive effects (Dale & Frank, 2017; Langellotto & Denno, 2004; Raupp et al., 2012; Bennett & Gratton, 2012; Burkman & Gardiner, 2014). For example, Parsons & Frank (2019) found that predators of the crape myrtle aphid *Tinocallis kahawaluokalani* were more abundant when the Shannon diversity of ground vegetation was higher. Greater availability of alternative prey and shelter from intraguild predation were mentioned as drivers of those observations. Parasitoids of *C. ohridella* in Europe are generalist parasitoids of leaf miners, attacking a wide range of hosts in various insect orders (Girardoz et al., 2007b). Although *C. ohridella* abundance or damage was not affected by tree diversity, parasitoids might have taken advantage of the presence of other insects on other tree species as supplementary or complementary prey resources. Additionally, a lack of synchronization between *C. ohridella* and its main parasitoids has been suggested (Grabenweger, 2003; Grabenweger, Avtzis, et al., 2005a). Overwintering parasitoids tend to emerge at the same time as the moth, several weeks before the presence of *C. ohridella* nymphs, their main hosts. If parasitoids emerge too early in the season, they might need to develop one generation on alternate host, such as leaf miners found on maple or other trees species (Grabenweger, 2004). As such, it has been suggested that parasitism rate of *C. ohridella* may be higher in more diverse environments, as observed in our study (Girardoz et al., 2006). However, and contrary to our prediction, we found no effect of the abundance of neighboring sycamore maple trees on parasitism rate, although it has been mentioned that parasitoids targeting leaf miner of the sycamore maple often shift to *C. ohridella* when its infestation are high (Girardoz, Volter, et al., 2007c).

Although more parasitized, horse chestnut leaf miners did not make lower damage in more diverse urban forest patches, which contradicts the associational resistance hypothesis. It has been mentioned that parasitism probably plays a minor role in the population dynamics of *C. orhidella* (Freise et al., 2002; Grabenweger, Avtzis, et al., 2005b). It is therefore likely that parasitism pressure was not the main driver of abundance and damage of an invasive pest already well established in the city. Finally, it is important to note that we focused only on parasitism in this study, but predation rate by birds or arthropods has also been documented for this species (Grabenweger, Kehrli, et al., 2005b).

## Conclusion

In conclusion, we showed that tree neighborhood affected the abundance and damage of a non-native invasive pest in an urban environment. Specifically, we showed that tree density, more than tree diversity, affects leaf insect damage by *C. ohridella* while tree diversity enhanced the parasitism rate of the exotic pest. This study, while not providing additional evidence of associational resistance, suggests a greater effect of urban tree diversity on top down processes (regulation by natural enemies) than on bottom up processes (accessibility of tree resources) in cities. The specific dispersal and colonization processes of *C. ohridella* in urban environments and the management of leaf litter in cities may have masked the direct effects of resource concentration. In the current context of increasing invasions of non-native pests in urban forests, additional studies appear necessary at a larger spatial scale, taking into account not only the composition but also the fragmentation and connectivity of tree patches to better predict and prevent the risk of damage.

## Acknowledgments

We thanks Tom Stemmelen for his help designing the Figure 1. This study was conducted in the framework of the HOMED project, which received funding from the European Union’s Horizon 2020 research and innovation program under grant <u>771271</u>.

